# Birds are easier to trick: an effect of magnetic field manipulation on migratory orientation of Nathusius’ pipistrelle in the circular release box

**DOI:** 10.1101/2023.02.20.529207

**Authors:** Fyodor Cellarius, Gleb Utvenko, Mikhail Markovets, Alexander Pakhomov

## Abstract

Bats, like birds, are capable of long annual migrations; however, a very limited number of studies are devoted to the role of the Earth’s magnetic field in bat navigation. We aimed to perform a series of experiments on Nathusius’ pipistrelle (*Pipistrellus nathusii*) to ensure that they are able to use the geomagnetic field for orientation. Bats were tested under two different conditions: in the geomagnetic field and the field, rotated 120° CW. To determine the takeoff direction and analyse behaviour in different magnetic conditions, we used the modified circular release box (CRBox) and a mini camera with IR LEDs. Helmholtz magnetic coils were used to manipulate the magnetic field. Bats were captured during migration through the Curonian spit (Kaliningrad region, Russia). Totally 53 bats were tested during August and September 2021-2022. During the second year, we recorded post-release bats’ behaviour using a thermal camera. Although results from 2021 are ambiguous, data obtained in 2022 suggests that under given conditions bats, unlike birds, could prefer local audible landmarks and wind direction prior to global cues. However, the recordings of released bats clearly show that they have some specific directional preferences, which correspond to their migratory orientation.

## Introduction

Every year, millions of animals migrate in search of food, shelter, and mating opportunities. Migration is well-developed in different classes of vertebrates and invertebrates as well: insects (Menz et al., 2022; Stefanescu et al., 2016), fish (Béguer-Pon et al., 2015), amphibian (Shakhparonov and Ogurtsov, 2016; Shakhparonov et al., 2022), reptiles (Luschi et al., 1998), birds (Salomonsen, 1967), and mammals can perform long-distant annual movements. In mammals, annual seasonal migrations can be performed by bats (Alcalde et al., 2021; Hutterer et al., 2005; Steffens et al., 2004; Vasenkov et al., 2022), whose navigational abilities even today remain extremely poorly understood in comparison with birds. Similarly impressive feats are found in humpback whales (*Megaptera novaeangliae*) on their 6500 km migration, which they can often complete with better than 1° precision, despite the effects of variable sea-surface currents (Horton et al., 2011).

Performing such migrations, animals can use a variety of information sources, including global astronomical and magnetic cues. The sensory basis of migratory navigation and orientation is best studied for passerine birds, thanks to the existence of a behavioral paradigm – Emlen funnels, that allow researchers to set up experiments on birds’ migratory behaviour under controlled conditions (Emlen and Emlen, 1966). Such a method existed for decades for birds; similar techniques were invented for butterflies and sea turtles (Lohmann, 1991; Mouritsen and Frost, 2002). Until recently, there was no similar behavioral paradigm and method for studying bats under controlled conditions. The invention of the circular release box (CRBox) for bats provided a powerful tool for understanding the sensory basis of bat orientation and navigation during migration (Lindecke et al., 2019a).

Thanks to the existence of such a controlled approach, a set of direct evidence for using a magnetic compass exists in birds (Wiltschko and Wiltschko, 1972). However, only indirect ones exist for bats. At the moment, the magnetic orientation and navigation of bats is a practically unexplored topic in chiropterology. Until recently, the only method available to researchers for studying the orientation and navigation of bats was their release with radio transmitters (Holland, 2007; Holland et al., 2008; Lindecke et al., 2015). Despite several obvious benefits, this method, however, does not allow scientists to manipulate the sources of navigational information available to bats (stars, the sun, and the magnetic field) during their migratory flight.

Currently, the majority of the studies are devoted to the analysis of the impact of strong magnetic pulses on bat orientation (Holland et al., 2006; Holland et al., 2008) and the hierarchy between sources of orientational information (magnetic field, stars, and the Sun). Thus, it was shown that greater mouse-eared bats *Myotis myotis* exposed at sunset to a changed magnetic field (rotated 90° clockwise) prior to release, alter their homing orientation relative to the control bats, exposed to the natural magnetic field (Holland et al., 2010). This indicates a calibration of the magnetic compass to the Sun, similar to what is shown in some migratory birds (Muheim et al., 2006), but unlike in birds, this process does not necessarily require access to polarized light, only access to the sun at sunset is important (Lindecke et al., 2015), but see (Greif et al., 2014). At the same time, only adult soprano pipistrelles *Pipistrellus pygmaeus*, exposed to the mirrored sunset before testing, show the orientation response; first-year subadults are not able to orient neither being exposed to natural nor mirrored sunset (Lindecke et al., 2019b). This may indicate that subadults fly with adults during their first migration, like in some bird species (Chernetsov et al., 2004).

The only attempts to measure directly the effect of magnetic field manipulation on bats’ behavior were performed on a Chinese noctule (*Nyctalus plancyi*). However, these bats were not in their migratory state and the effect was measured by analysing the bats’ roost locations inside a small experimental box (Wang et al., 2007). Therefore, we aimed to find direct evidence of magnetic compass in migratory bats (Nathusius’ pipistrelle *Pipistrellus nathusii*), using a recently proposed CRBox approach.

## Materials and Methods

### Model species and study site

Nathusius’ pipistrelle *Pipistrellus nathusii* was chosen as a model species for the present study. This species is a long-distance bat which can perform over 2000 km migration each year (Alcalde et al., 2021; Vasenkov et al., 2022) and one of the typical model animals to study orientation and navigation in migratory bats (Lindecke et al., 2015; Lindecke et al., 2021). All bats (male and female, adults) were captured on the Curonian Spit (Kaliningrad region, Russia; **Figure S1**) using a custom-made telescoping high mist-net bat system during their autumn migration from the middle of August to the end of September in 2021-2022. After capture, bats were aged based on the closure of the epiphyseal gaps (additionally, we took the photo of this part of a bat’s wing for further rechecking), weighted (for experiments we used only bats with seasonally appropriate body mass ≥ 7.0 g, according to (Lindecke et al., 2021; Voigt et al., 2012) and ringed. The current research was carried out in compliance with the ARRIVE guidelines (https://arriveguidelines.org) and all procedures were approved by the Ethics Committee of the Zoological Institute, Russian Academy of Sciences (permit #5-15/10-08-2021). The bats were released directly back into the wild after tests.

### Experimental setup and test procedure

For condition-controlled experiments and analysis of bat behaviour and orientation in different magnetic conditions, we used a modified version of a circular release box (CRBox; (Lindecke et al., 2019b)). Our experimental setup consists of two parts: a circular arena (40 cm diameter, **4** in **Figure 1**) and a plastic cylinder (42 cm diameter, 54 cm height, **1** in **Figure 1**). We used OpenSCAD (Kintel and Wolf, 2014) open-source software to create a 3D model of the arena, printed it (PETG plastic) and coated it with synthetic leather which supports the crawling of bats compared to plastic. The cylinder is made from thin PVC (2 mm) and attached to two circular wooden frames at the top and bottom. We covered the interior wall of the cylinder with a uniform black felt to reduce the reflection from IR LEDs. A transparent circular plexiglass lid (diameter 58 cm, **5** in **Figure 1**) with an 11 cm hole in the centre plays the role of a roof of the CRBox. The lid extends 9 cm beyond the arena, and this part of the lid is covered with black paint, which created a uniform environment inside the CRBox and prevented the bats from seeing the stars and the night sky overhead when they reached the edge of the arena (in 2021 there was no transparent lid, only a 6 cm plastic circular hoop that hinders the bat from seeing the night sky similar to the lid in 2022). The upper part of the cylinder is covered with a white acrylic lid (60% light transmission; **9** in **Figure 1**) and is equipped with a remote video recording and release system (**6** in **Figure 1**) placed inside a grounded aluminium box. This system consists of two parts: a microcontroller Arduino Uno with a stepper motor (**7** in **Figure 1**) and a small single-board computer Raspberry Pi 4, which controls both a wide-angle miniature IR camera (**8** in **Figure 1**) and motor rotation via custom-written script. Raspberry PI is connected to a PC tablet via FTP cable (30 meters in 2021 and 50 meters in 2022) for real-time observation of bat’s behaviour inside the CRBox and running the script.

**Figure 1.**
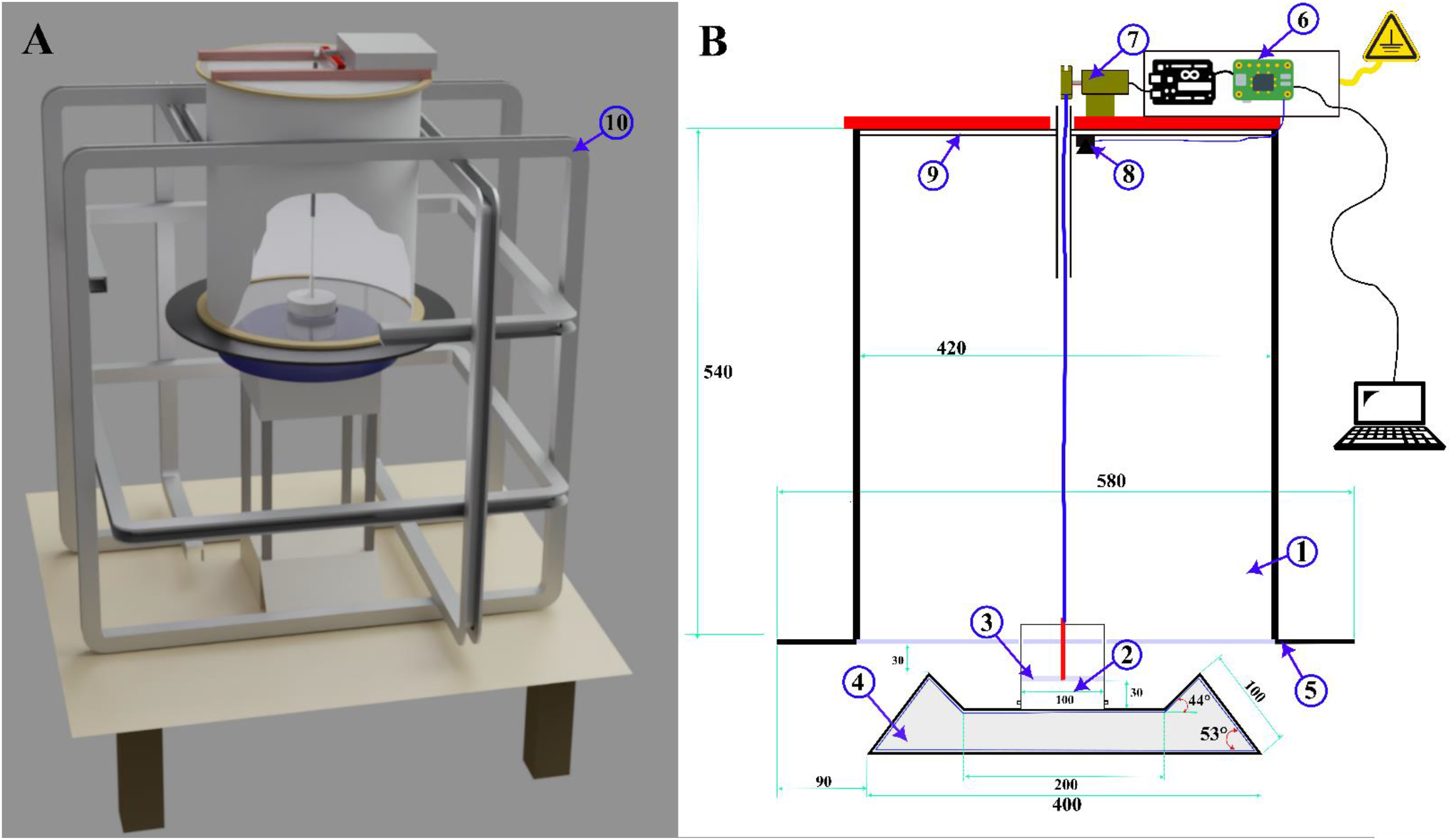
Experimental setup. A) a 3d model of whole setup placed at the release site; B) the drawing of a modified version of the CRBox used in our experiments. 1 – the plastic PVC cylinder, 2 – the acclimatization box, 3 – a transparent lid of the box, 4 – the main arena of the CRBox, 5 – transparent Plexiglass lid (a “roof” of the arena), 6 - an aluminium box with Raspberry Pi 4 and Arduino, 7 – a stepper motor, 8 – a miniature video camera, 9 - a white acrylic lid, 10 – the magnetic coils.

Experimental magnetic fields were produced by three-dimensional Helmholtz coils (1 m diameter, ‘magnetic coils’ hereafter; **10** in **Figure 1**). This coil system was identical to the one used by the Rybachy animal navigation group in previous behavioural experiments (Chernetsov et al., 2011; Pakhomov and Chernetsov, 2014). We used a 12V car battery with a custom-made control box placed in a shielded and grounded aluminium box to power the magnetic coils. Using car batteries instead of typical power supplies in our previous studies (Kishkinev et al., 2015; Pakhomov et al., 2022) allows us to exclude artificial sound produced by supplies as a factor that could affect the takeoff direction of bats from the release setup. The magnetic coils were placed on the wooden table (50 cm height); thus, the CRBox inside the coils was about 1 m above ground. Magnetic parameters were checked and calibrated before each magnetic condition (the natural magnetic field or NMF and a 120° counterclockwise changed magnetic field or CMF) using a portable magnetometer (FMV400 model, MEDA Inc., USA).

After capture and ringing, pipistrelles were immediately transferred to the experimental release site (RS 3 in 2021 or RS 4 in 2022, **Figure S1**) if weather conditions were favourable. If we had no opportunity to test them after capture, we transferred bats to a wooden house (a cool, dark, and quiet environment) to keep them in groups of 5-9 individuals in plastic boxes and test the following night. The bats were provided with fresh water and a mixture of blended black soldier fly larvae and mealworms using a syringe in the evening before nocturnal tests (the weight of each bat after capture and feeding was similar). In contrast to previous studies (Lindecke et al., 2019a; Lindecke et al., 2019b), we had no opportunity to translocate captured bats far away from the seashore and capture sites during tests due to geographical features of the Curonian Spit (Figure S1). All bats were kept inside a plastic bucket during the test night without seeing the local surroundings, at least 30 m from the release site. In 2022 wind direction and speed, and air temperature at the release site were measured using a portable anemometer (CEM DT-618, Shenzhen Everbest, China), in 2021 we measured only speed and temperature.

Each test consisted of two phases: acclimatization and release. During the first phase, we randomly chose a bat from the bucket and placed it into the acclimatization box (10 cm diameter; **2** in **Figure 1**) with a transparent top lid (**3** in **Figure 1)** to observe bat behaviour inside the box. The direction of insertion of bats into the box was changed randomly (by 90° between trials) and bats did not see any signs of the local environment during translocation from the keeping site to the release site and insertion. We waited for 10 s after this procedure and started to record a video of bat behaviour during the first phase. After 5 min of acclimatization, the script automatically stopped video recording (+ saving a file to SD card), started recording a new video and lifted the box to release the bat simultaneously. The release phase lasted 5 min and if the bat did not leave the CRBox during this time, a trial was canceled. Before the next trial, the arena was completely cleaned with 70% ethanol to remove any olfactory cues from previous bats. There were several NMF and CMF test blocks during each experimental night: we released 5 bats in NMF (control), then the next 5 bats in CMF (experimental), etc. Additionally, in 2022 we recorded tracks of free-flying bats after leaving of the CRBox (see **Video S1**) using a thermal monocular (xEye E3W model, IRay Technology Co., Ltd., China) and used a bat detector (Petterson D320, Pettersson Electronik, Sweden) to control the presence of conspecific bat calls at the RS. All untested bats were released unharmed in the wild at the end of the experimental night.

### Data analysis and statistics

We determined the bats’ orientation along with takeoff latency (time necessary for a bat to depart, in seconds) in the CRBox based on the release phase video. We used two measures of orientation for a single bat: crawling direction (direction to the point where a bat reached the highest part of the arena; **1** in **Figure S2**) and takeoff direction (direction in which a bat emerges from the arena; **2** in **Figure S2**). We used the Rayleigh test (Batschelet, 1981) as a criterion for the unimodality of samples; the same test was used for doubled angles in the case of bimodal samples (Batschelet, 1981). To test for significant differences in distribution and, in particular, concentration in bats’ orientation under control and test conditions, we used a non-parametric Mardia-Watson-Wheeler (MWW) test (Batschelset, 1981) and Concentration test (Mardia and Jupp, 2000), respectively. Nonparametric Mann-Whitney U-test was used to compare linear data such as wind speed and takeoff latency. All tests were performed using python 3.9 (Van Rossum and Drake, 2009).

## Results and Discussion

Totally, 53 adult bats in good condition (mean body mass 9.2 ± 1.1 g) were released from our experimental setup in 2021-2022. The observed crawling and takeoff directions are approximately the same for all animals, regardless of the experimental conditions (**Figure S3**; see raw data in **Table S1** for details). Therefore, further discussion will be focused on the takeoff directions, similar to previous studies on Nathusius’ pipistrelle (Lindecke et al., 2019b).

In 2022, bats did not show a significant unimodal orientation relative to the geographic north, neither in the control nor in the experimental group: α = 36°, r = 0.32, p = 0.23, n = 14, (**Figure 1A)** and α = 193°, r = 0.22, p = 0.59, n = 11 (**Figure 1B**), respectively. However, the animals in NMF showed a bimodal orientation corresponding to an axis perpendicular to the spit (results for doubled angles: α = 109°/289°, r = 0.56, p = 0.01, n = 14), in contrast to the bats that were released in CMF (doubled angles: r = 0.23, p = 0.55, n = 11). In addition to the direction relative to the geographic north, we analysed takeoff directions normalized to the wind source direction. In the control group, the animals emerged the arena mainly against the wind (α = 24°, r = 0.52, p = 0.02, n = 14; **Figure 1C**), while the experimental animals showed no significant orientation (α = 99°, r = 0.25, p = 0.51, n = 11; **Figure 1D**).

In 2021, pipistrelles were not significantly oriented (unimodal or bimodal) in either the experimental (α = 109°, r = 0.24, p = 0.43, n = 15; doubled angles: r = 0.33, p = 0.19; **Figure S4A**) or control group (α = 179°, r = 0.39, p = 0.14, n = 13; doubled angles: r = 0.33, p = 0.23; **Figure S4B**). If we consider reliable bats with weight ≥ 8 g, the control bats were oriented (bimodal), similar to 2022 (doubled angles: α = 142°/322°, r = 0.55, p = 0.03, n = 11; **Figure S4C**), and bats from the experimental group showed tendency to orient in southern direction (α = 180°, r = 0.50, p = 0.048, n = 12; **Figure S4D**). We did not record the wind direction in 2021, so it is not possible to analyse the effect of wind on takeoff direction. Additionally, results of our 2021 experiments cannot be considered reliable due to several reasons and cannot be used for further discussion: 1) the experimental setup in 2021 had no stoppers for the acclimatization box, therefore its wide movements could additionally stress the animals; 2) there was skyglow in southern direction due to the light pollution from the nearest town which was visible in RS 3 (2021) but not in RS 4 (2022) and could attract bats like birds (phototactic response to setting sun or the brightest part of the sky during tests in circular arenas, (Muheim and Jenni, 1999).

In our study, bats from the experimental group which were released in 120° CW deflected magnetic field were not oriented (unimodal or bimodal), in contrast to control bats. There are no significant differences in distribution or concentration between the control and experimental groups, neither in the case of comparing samples normalized to wind direction (MWW test: W = 1.67, p = 0.4; Concentration test: test score = 0.81, p = 0.4) nor in the case of samples normalized to geographical north (MWW: test score = 4.71, p = 0.1). Additionally, we did not find any differences in takeoff latency between the control and experimental groups (Mann-Whitney U test: U = 67.5, p = 0.6; **Figure S5**). However, despite the lack of death proof, we can assume that a dramatic change in the magnetic field disturbs the orientation of bats in a circular arena.

In contrast to previous studies in migratory bats (Lindecke et al., 2019a; Lindecke et al., 2021), Nathusius’ pipistrelles released from the CRBox did not show southwestern, typical for migratory animals (birds and butterflies) on Curonian spit (Chernetsov et al., 2020; Pakhomov et al., 2017; Pakhomov et al., 2022) (Pakhomov et al., in prep). There are several assumptions that can explain our results:

1) in the presence of two strong cues (wind direction and sea/lagoon noise), bats in NMF can choose takeoff direction in accordance with them. In the study with another version of the CRBox, Soprano’s bats have shown bimodal orientation towards the coastline and along their migratory route (Lindecke et al., 2019a). The authors proposed that some bats tried to return to the original migratory pathway using sea noise as a cue source. However, this assumption cannot be applied to our case because Nathusius’ bats were tested within their migration corridor which can cover the entire width of the spit, according to measurements of conspecific bat calls. Nathusius’ bats prefer a weak headwind during migration (Pētersons, 2004), which corresponds to flying against the wind from our setup. We assume that control bats implemented so-called taxon navigation (Redish, 1999), choosing takeoff direction in correspondence with two strong directional cues: sea/lagoon noise and wind directions.

2) Another important factor that may influence bats’ behaviour and takeoff direction is the presence of magnetic coils in their emergence way. Despite the magnetic coils are a symmetric object and have 8 symmetric windows (4 big E-W and N-S windows, 46 × 46 cm; 4 small radial or NE-SW and NW-SE windows, 35 × 46 cm), coils can be identified by bats as an obstacle. In such a hypothetical situation, firstly they can try to leave such a stressful place using environmental cues (wind or sea sound, for example) and then bats use other cue sources (stars or the magnetic field) to determine migratory direction. In 2022, we additionally recorded animals’ behaviour after takeoff with a thermal camera. Unfortunately, we can only operate with the projection of animal tracks and their vanishing bearings in the Northeast-Southwest plane. However, according to these data, the animals choose to a greater extent the Southwestern sector (n_Southwest_ = 23, n_Northeast_ = 8, Binominal test for P=0.5: p = 0.01; **Figure 2A**), which corresponds to their natural migratory direction. There are no such preferences were observed in takeoff orientation (n_Southwest_ = 10, n_Northeast_ = 15, Binominal test for P=0.5: p = 0.4; **Figure 2B, 2C**). The majority of the released bats showed circling behaviour after takeoff (see **Video S1**). The same behaviour was occasionally observed in the previous study (Lindecke et al., 2019a). However, this phenomenon was not a consistent pattern, unlike in our study: in 2022, a few released bats fled straight right after release, but most of them showed at least a few circular turns. These results correspond to the radio telemetry data (Lindecke et al., 2015), which shows that some bats spend time near the release site before departure. We were not able to track bats’ flying after release for a long time, however, some individuals, apparently, did the same. This circling behaviour can be interpreted as an attempt by animals to integrate information from different ‘compass’ or ‘map’ cue sources (Baird et al., 2012; Douglas, 2021).

**Figure 2.**
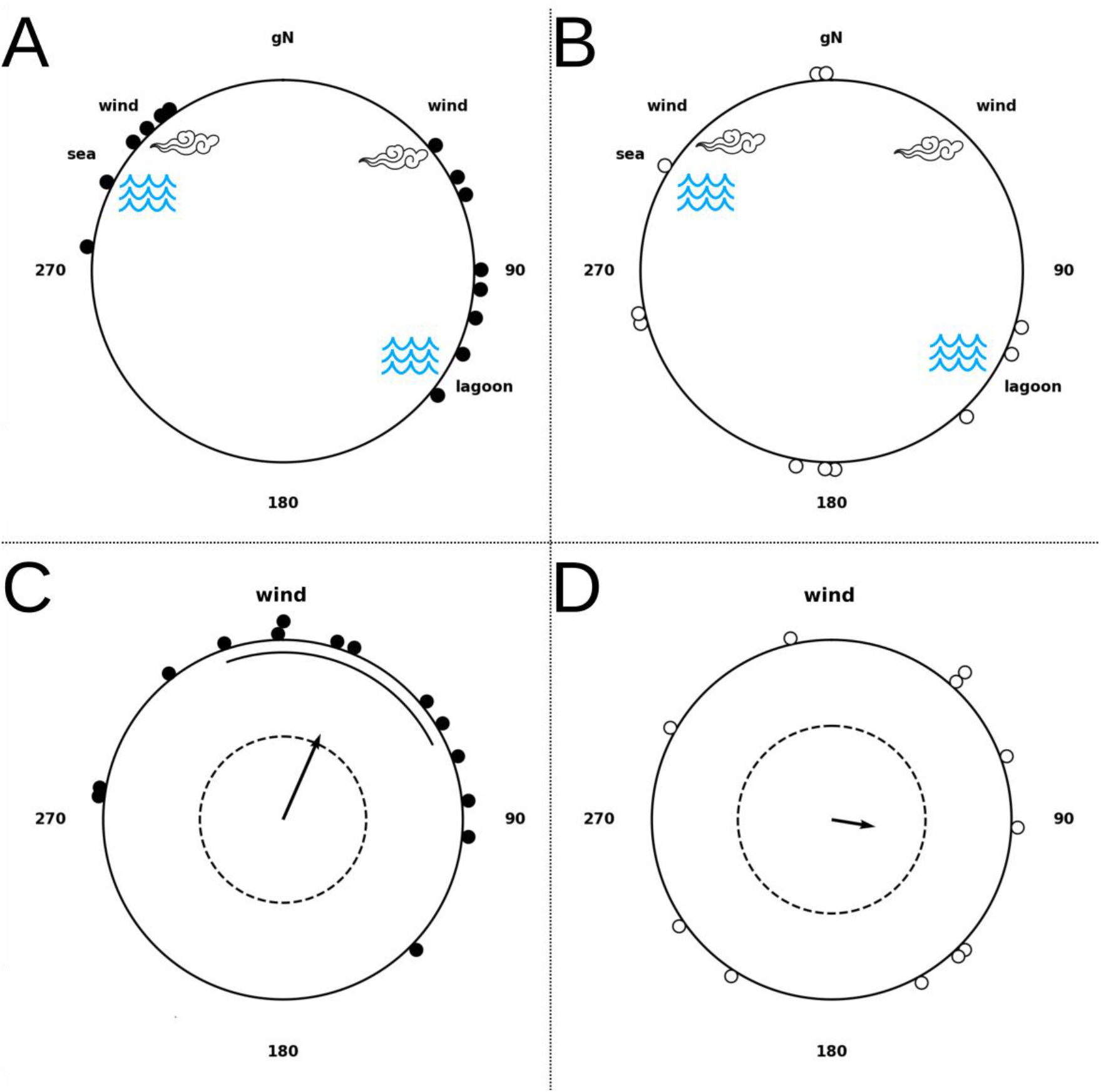
Orientation of Nathusius’ pipistrelles in 2022 under different experimental conditions relative to geographic north and wind: A) under natural magnetic field (NMF), relative to geographical north; B) under the field, rotated 120° CCW (changed magnetic field, CMF), relative to geographical north; C) under natural magnetic field, relative to the wind; D) under the field, rotated 120° CCW, relative to the wind. Each dot at the circle periphery indicates the orientation of one individual bat. Black dots correspond to animals under NMF, and white dots to animals under CMF. The inner dashed circle represents the 5 % significance level of the Rayleigh test. Geographic North (gN) corresponds to 0° at the top plots (A, B); wind source direction corresponds to 0° at the bottom plots (C, D). Wind source direction denoted on the upper plots as “wind”; directions towards the sea or lagoon as “sea” and “lagoon”, respectively. The inner arc on plot C indicates the 95 % confidence interval for the mean orientation direction.

**Figure 3.**
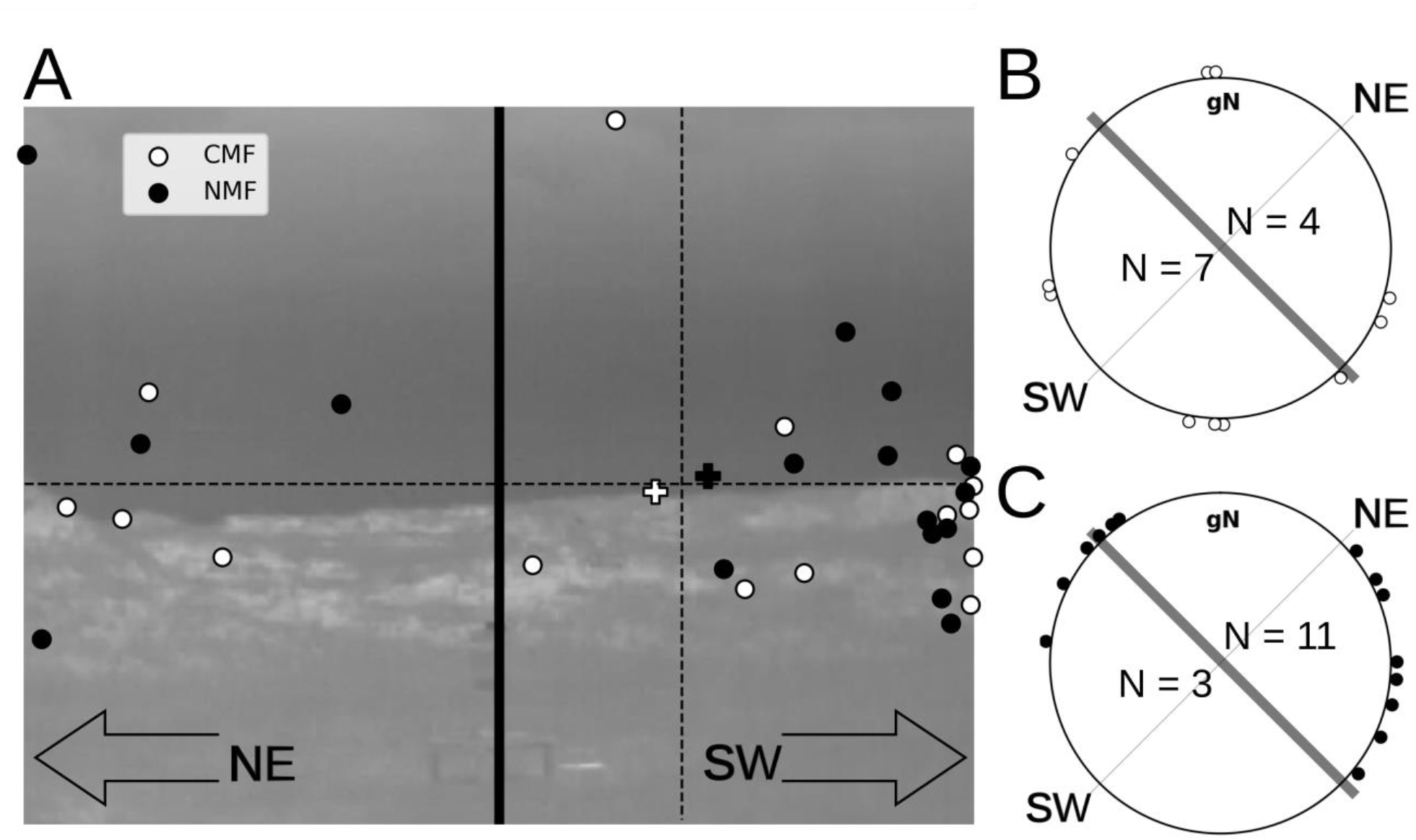
Projection of vanishing bearings of Nathusius’ pipistrelles in 2022 under different experimental conditions in the Northeast-Southwest plane, according to the thermal camera (A) and takeoff orientation of the bats in the circular arena (B, C). White dots on the thermal image represent the vanishing bearings of individual bats, released from a changed magnetic field (CMF); black dots correspond to bats released from a natural magnetic field. The same colors on circular diagrams correspond to the individual orientation of bats under CMF (B; white) and NMF (C; black), respectively. The bold black line on the thermal image and bold gray lines on circular diagrams represent the border between the northeast and southwest sectors. The bold black line crosses the centre of our experimental setup (the magnetic coils and the CRBox). White and black “plus” markers on the thermal image show centres of mass for vanishing bearings of bats, released from CMF and NMF, respectively. The intersection of dashed lines shows the centre of mass for all animals.

Soprano pipistrelles tested in the classical version of the CRBox at Pape Biological Station (PBS), Latvia (Lindecke et al., 2019b) were able to choose an appropriate migratory direction, in contrast to Nathusius’ pipistrelle released from the modified version of the CRBox at Biological Station Rybachy (BSR) on the Curonian Spit. There are several key differences between these release sites (BSR and PBS): clearly audible landmarks (the lagoon and the sea) and migratory corridor due to the width of the Curonian spit and influence of the wind speed and direction. In our case, the wind was significantly stronger than in the study where the method used was first proposed (Lindecke et al., 2019a); Mann-Whitney U-test: z = 1248, p << 0.01; **Figure S6**). Despite the wind and strong sound cues, this experimental site is well suited for similar orientation experiments on migratory birds (Kishkinev et al., 2013; Kishkinev et al., 2015; Pakhomov et al., 2018). It might be possible due to the fundamental difference between Emlen funnels and CRBox: Emlen funnels exploit an accumulation effect since a bird is locked inside the funnel and remains active for a long time and usually is tested several times before releasing. Additionally, in 2020 Nathusius’ bats in our preliminary tests performed in the classical version of the CRBox showed the tendency in southwestern direction (α = 227°, r = 0.49, p = 0.12, n = 9). Together with data from the thermal camera, results of all our orientation experiments on migratory bats in 2020-2022 might indicate that migratory bats are not so easier to trick if you are trying to test them under the same experimental conditions as migratory birds and other migratory animals.

## Supporting information

Video from the thermal camera

## Acknowledgements

We are grateful to Nadezhda Romanova and Inga Iokhina for assistance in the creation of experimental setups, bat catching and experiments.

## Authors’ contributions

A.P. and F.C. designed research; G.U., F.C. and M.M. caught and ringed bats; A.P., F.C. and G.U. performed experiments, collected and analysed the data; A.P., F.C. and G.U. wrote the first draft of the manuscript. All authors commented on the manuscript and gave final approval for publication.

## Data Availability

All data generated or analysed during this study are included in this manuscript (and its Supplementary Information files) or available from the corresponding author on reasonable request.

