## Supplementary materials (Figures) for "Birds are easier to trick: an effect of magnetic field manipulation on migratory orientation of Nathusius’ pipistrelle in the circular release box"

**Figure S1. Location of capture and release sites on Curonian Spit, Russia.**

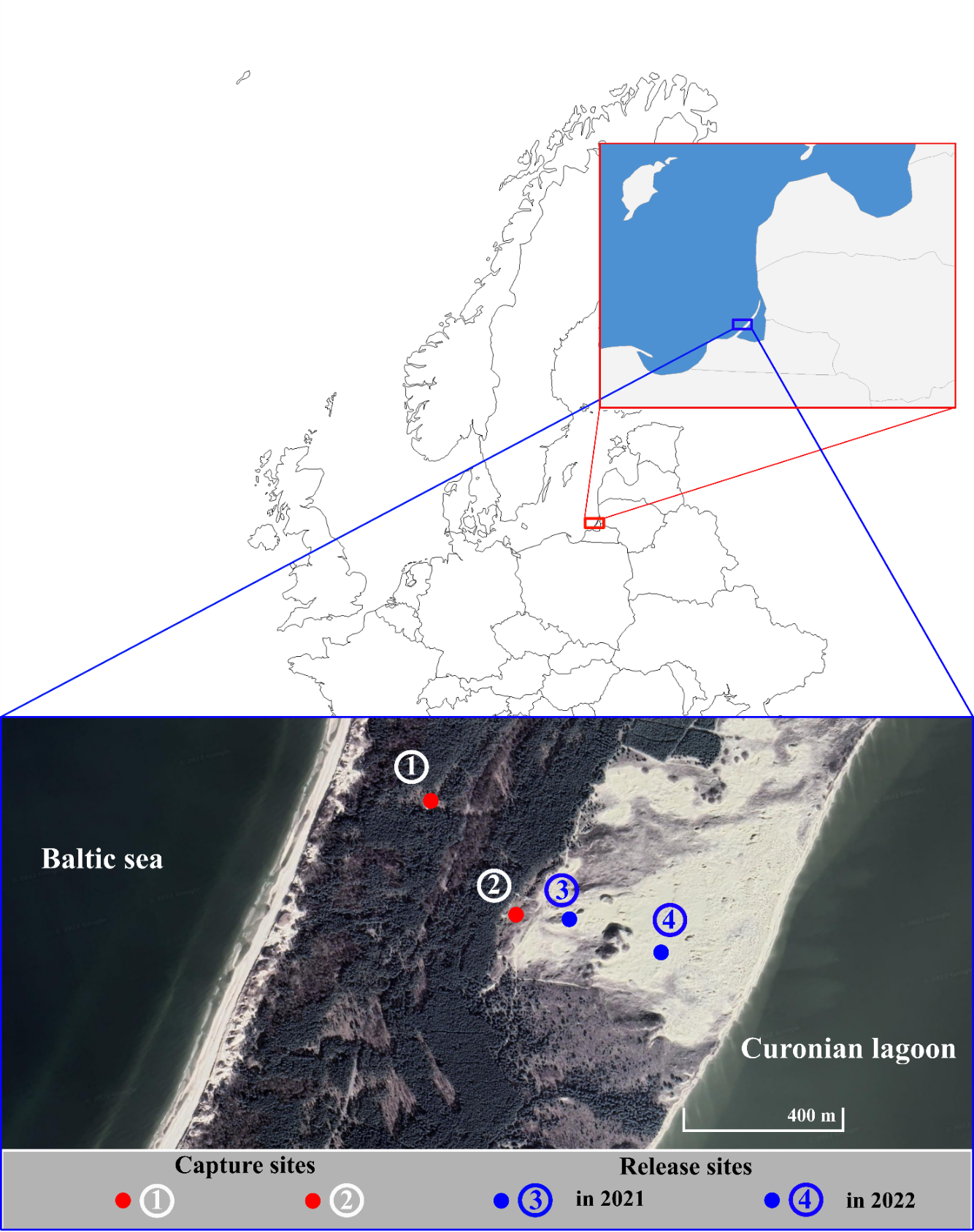

**Figure S2. Frame from the video after lifting the acclimatization box.** The parameters determined from such a video recording are: departure time, departure direction and crawl direction (for more details, see the text). The yellow circle is an isoline showing the greatest height of the arena walls, the bat is highlighted with a red rectangle. The blue line symbolizes the hypothetical route the animal followed when leaving the arena. Two stars on the route are the points by which the crawl direction (1) and the takeoff direction (2) were determined (they could not coincide). The direction of departure was determined by the point from which the bat left the arena, and the direction of crawling out was determined by the point where the animal crossed the inner boundary of the arena (yellow circle).

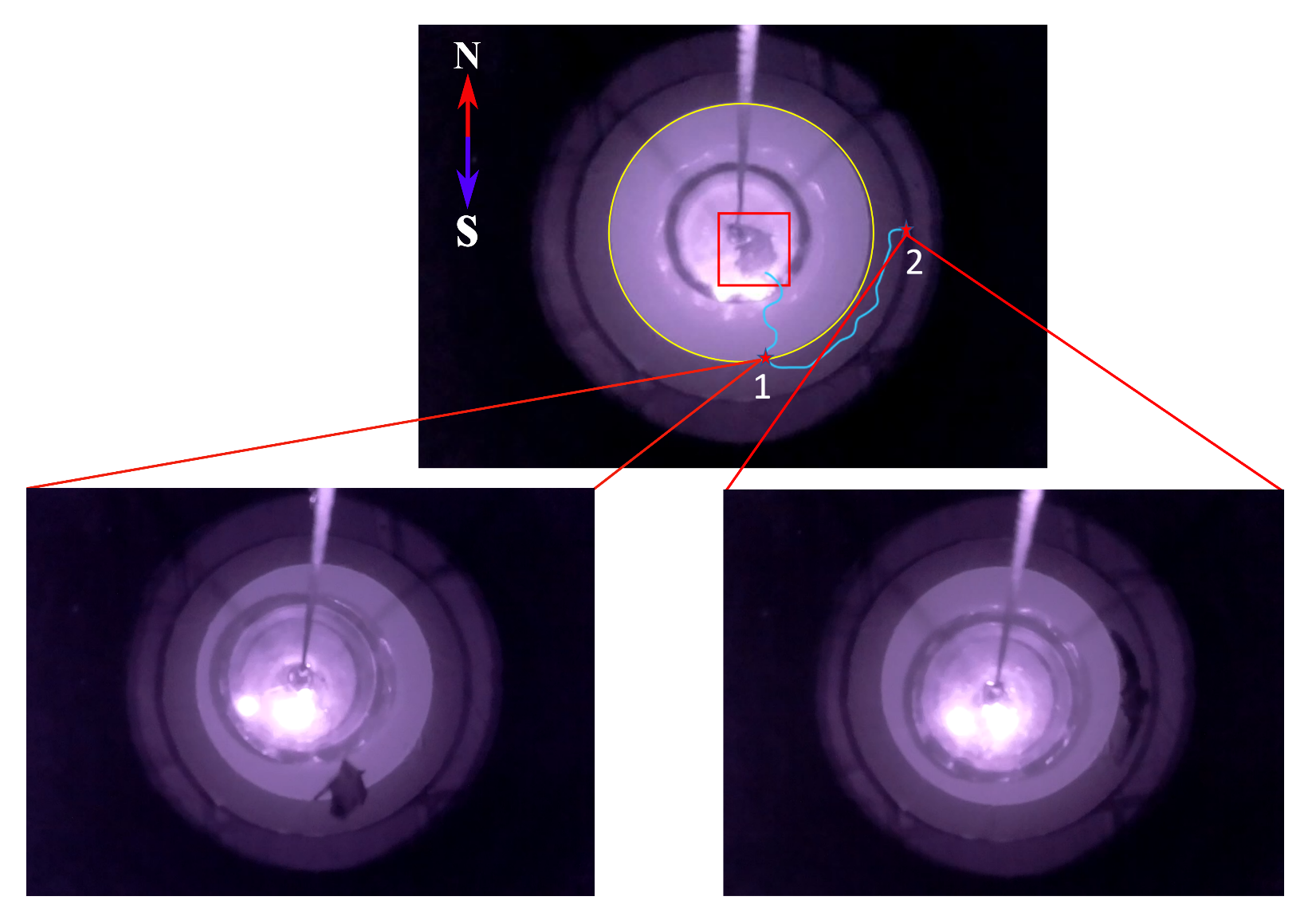

**Figure S3. Takeoff orientation of Nathusius’ pipistrelles in 2022 normalized on their crawl orientation.**

Each dot at the circle periphery indicates the orientation of one individual bat. Black dots correspond to animals under NMF, and white dots to animals under CMF. The inner dashed circle represents the 0.001 significance level of the Rayleigh test.

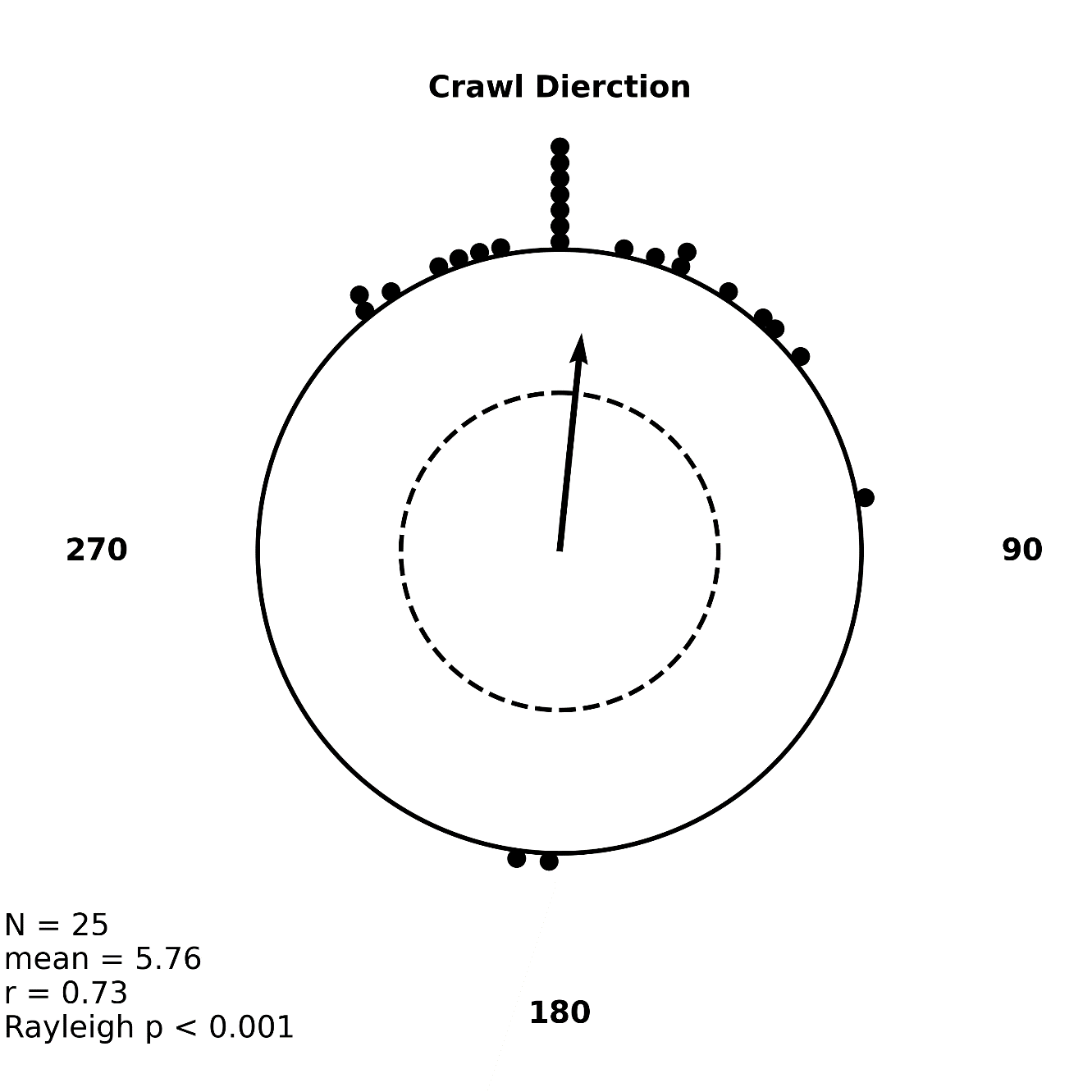

**Figure S4. Orientation of Nathusius’ pipistrelles in 2021 under different experimental conditions relative to geographic north**: A) under natural magnetic field (NMF); B) under the field, rotated 120° CCW (changed magnetic field, CMF); C) only bats with weight >= 8 g, NMF; D) only bats with weight >= 8 g, CMF. Each dot at the circle periphery indicates the orientation of one individual bat. Black dots correspond to animals under NMF, and white dots to animals under CMF. The inner dashed circle represents the 5 % signiﬁcance level of the Rayleigh test. Geographic North (gN) corresponds to 0°. Directions towards the sea or lagoon are denoted as “sea” and “lagoon”, respectively.

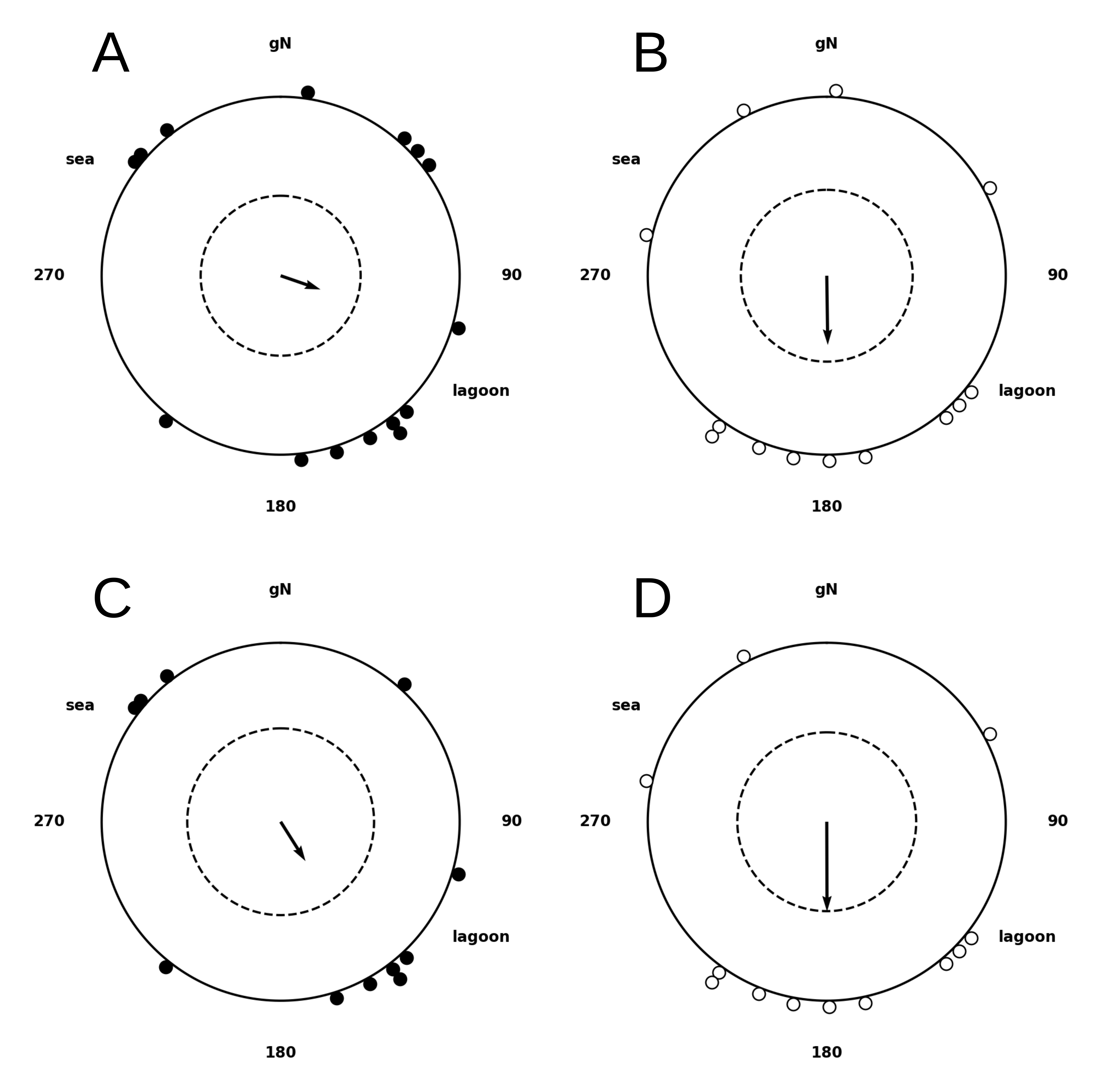

**Figure S5. Takeoff latencies (time between acclimatization box lifting and bat’s takeoff, in seconds) for animals under natural magnetic field (NMF) and field, rotated 120° CCW (changed magnetic field, CMF).**

Each dot corresponds to an individual bat’s latency. Whiskers show sample ranges, boxes represent Q1 and Q3, and the median is shown as an orange line.

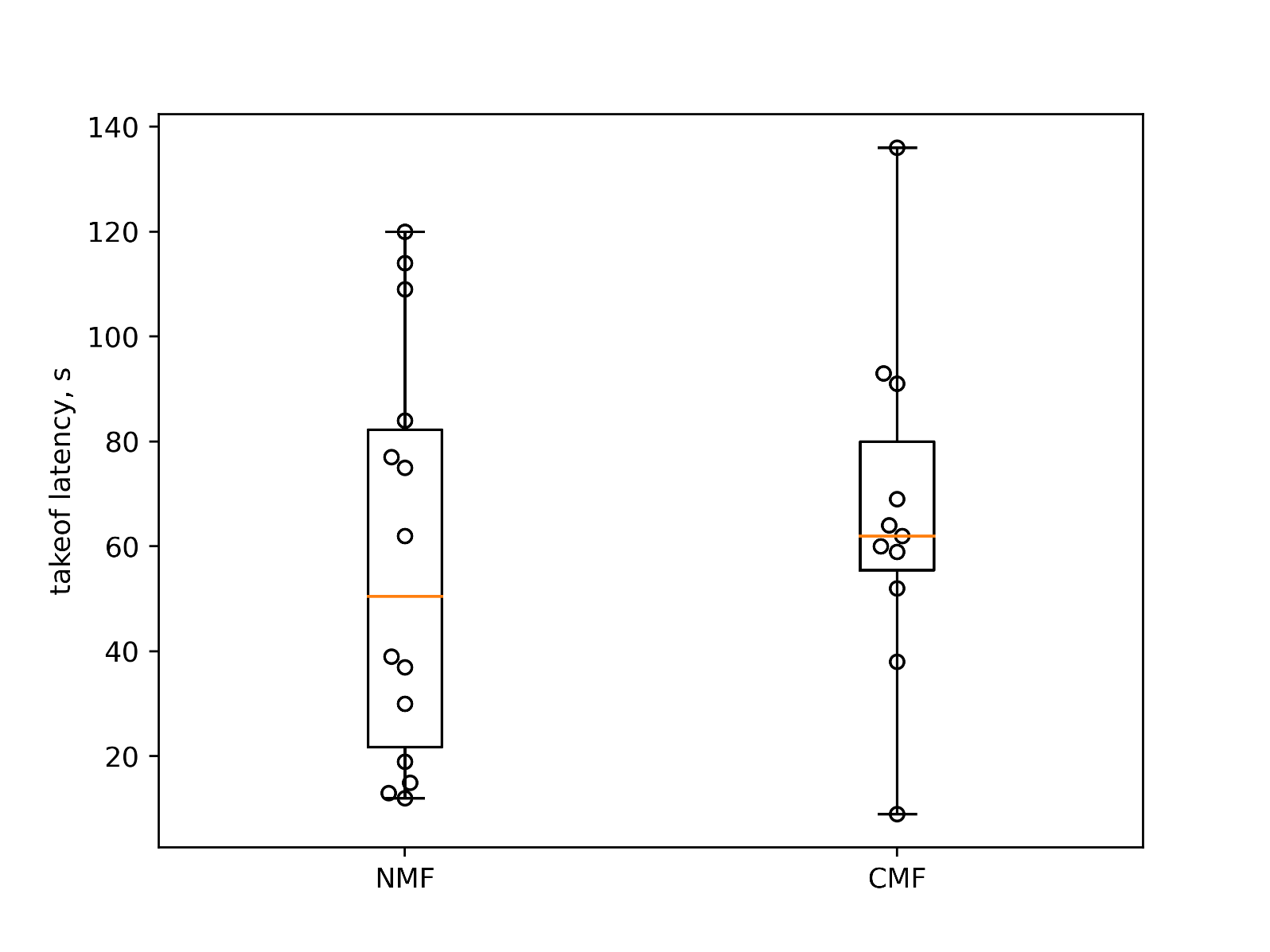

**Figure S6.** Combined violin and swarm plots of wind speed at our experimental site in 2022 and the corresponding data from Lindecke et al., 2019b. Each dot represents the wind speed corresponding to an individual bat’s takeoff. Curved lines show an estimated probability density for a wind speed.

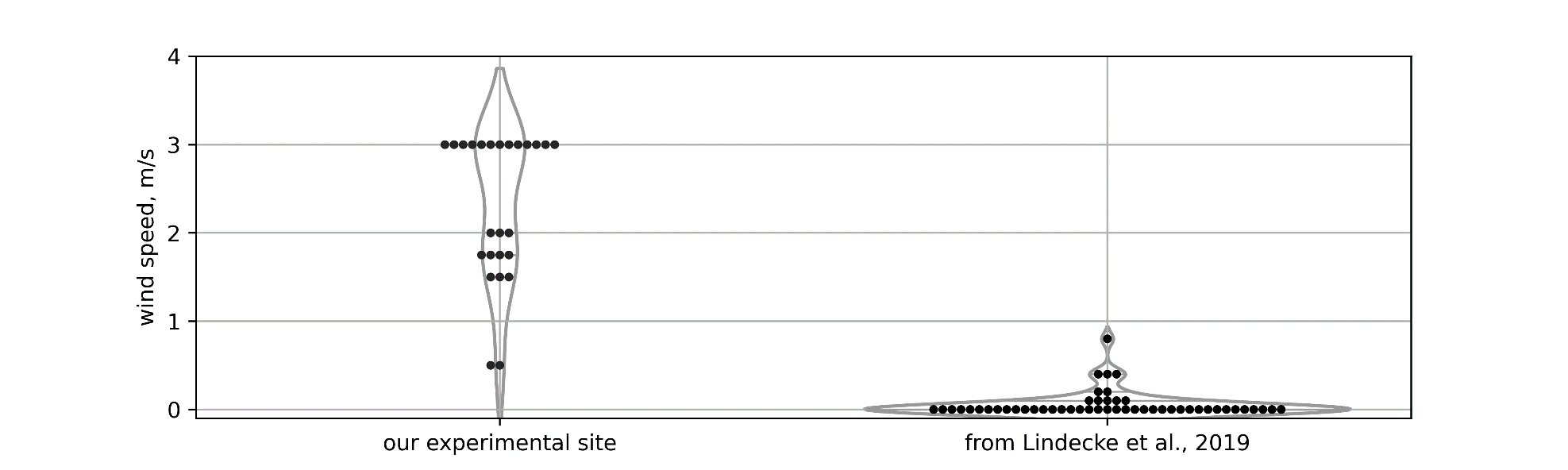
